# Sustained antibody response to ZIKV infection induced by NS1 protein is accompanied by the progressive appearance of autoreactive antibodies and cross-reactive B cell clones

**DOI:** 10.1101/2020.12.24.423863

**Authors:** Cecilia B. Cavazzoni, Vicente B. T. Bozza, Lucas Tostes, Bruno Maia, Luka Mesin, Ariën Schiepers, Jonatan Ersching, Romulo L.S. Neris, Jonas N. Conde, Diego R. Coelho, Luciana Conde, Heitor A. de Paula Neto, Tulio M. Lima, Renata G.F. Alvim, Leda R. Castilho, Ronaldo Mohana-Borges, Iranaia Assunção-Miranda, Alberto Nobrega, Gabriel D. Victora, Andre M. Vale

## Abstract

Besides antigen-specific responses to viral antigens, humoral immune response in virus infection can generate polyreactive and autoreactive antibodies. Dengue and Zika virus infections have been linked to antibody-mediated autoimmune disorders including Guillain-Barrè syndrome. A unique feature of flaviviruses is the secretion of non-structural protein 1 (NS1) by infected cells. NS1 is highly immunogenic and antibodies targeting NS1 can have both protective and pathogenic roles. In the present study, we investigated the humoral immune response to Zika virus NS1 and found NS1 to be an immunodominant viral antigen associated with the presence of autoreactive antibodies. Through single B cell cultures, we coupled binding assays and BCR sequencing, confirming the immunodominance of NS1. Of note, we demonstrate the presence of self-reactive clones in germinal centers after both infection and immunization, some of which clones presenting cross-reactivity with NS1. Sequence analysis of anti-NS1 B cell clones showed sequence features associated with pathogenic autoreactive antibodies. Our findings demonstrate NS1 immunodominance at the cellular level as well as a potential role for NS1 in ZIKV associated autoimmune manifestations.

## Introduction

The protective function of antibodies is instrumental for the control of most viral infections. However, in addition to antigen-specific responses to viral antigens, it has long been noted that viral infection can be accompanied by the appearance of polyreactive and often autoreactive antibodies of unknown function [1]. Emergence of self-reactive immunoglobulins has been reported for multiple viral diseases; these autoreactive antibodies have the potential to lead to autoimmune manifestations, which can be transient or long-lasting (reviewed in [2]). The origin of the stimulus driving autoantibody generation in viral infection remains controversial. On one hand, antigen mimicry between viral and self-antigens may explain the appearance of selected autoantibodies. Alternatively, a role for cytokine storm leading to non-specific, polyclonal B cell activation followed by disruption of B cell repertoire homeostasis and self-tolerance has also been proposed [3].

Recently, dengue virus (DENV) and zika virus (ZIKV) infections have been linked to the occurrence of autoimmune disorders of vascular, ophthalmic or neurological origin, including Guillain-Barrè syndrome [4, 5], in which autoantibodies seem to play a prominent role [6]. A unique feature of DENV, ZIKV and other flaviviruses is the abundant secretion of the non-structural protein 1 (NS1) in its hexameric form by infected cells [7-10]. While intracellular NS1 was shown to be necessary for viral replication, its role as an extracellular soluble factor is poorly understood [11]. It has recently been shown that NS1 can lead to endothelial dysfunction [12]. Moreover, work from multiple groups has shown that NS1 is highly immunogenic [13-15]. Notably, antibodies to DENV NS1 can attenuate the outcome of severe dengue and vaccination with ZIKV and DENV NS1 have been shown to be protective in animal models [16-19]. These studies suggest a role for NS1 in the pathogenesis of flavivirus infections, as well as a protective role for anti-NS1 antibodies. However, anti-DENV NS1 antibodies have also been implicated in autoreactivity and may contribute to dengue pathology, possibly through antigen mimicry between NS1 and components of self [20-23]; these observations raise concerns about the safety of NS1 as a vaccine antigen, and further studies are necessary to understand the humoral immune response to this molecule.

Studies of the pathogenesis of human viral infections in animal models face several limitations due to innate viral resistance of mouse species to many human viruses. For this reason, multiple groups have developed models that rely on immunocompromised mice, such as IFNAR-deficient strains, which show greater susceptibility to viral infections [24-27]. Although such strategies are useful in studies of viral pathology, they are less useful for the analysis of humoral immune responses, as type-I interferons broadly influence acquired immunity and directly impact activation of B cells by modulating B cell receptor (BCR) signaling [28, 29] possibly also affecting clonal selection and entry into germinal centers (GCs) [30]. For these reasons, an immunocompetent mouse model of ZIKV infection is preferred for the study of antibody response. Accordingly, recent studies have used such models to characterize the T cell [31-33] and neutralizing antibody response to ZIKV [34]. Using immunocompetent mouse models, we and others have shown that CD4+ T cells activate a robust IFNγ-dependent B cell response, which is associated with production of neutralizing IgG2a antibodies that bind to ZIKV envelope proteins including envelope protein domain III (EDIII) and have been associated with virus neutralization. Of note, passive transfer of serum from infected immunocompetent A129 mice protected immunocompromised mice against lethal heterologous challenge with ZIKV [35-37].

In the present study, we investigated the humoral immune response to ZIKV NS1 using an immunocompetent mouse model of ZIKV infection as well as NS1 immunization. The antibody response in infected animals showed that NS1 is an immunodominant viral antigen. Importantly, humoral response to NS1 is associated with the presence of autoreactive antibodies, both after infection or immunization. In-depth analysis of B cell clone selection into GCs, coupling single B cell cultures with BCR sequencing confirmed the strong immunodominance of NS1 and revealed the presence of frequent self-reactive clones among GC B cells, some of which were cross-reacted with NS1. Anti-NS1 B cell clones were enriched in charged amino acid residues in the CDR-H3 region, a feature shared by self-reactive clones [38, 39]. Anti-NS1 clones also showed low levels of somatic hypermutation (SHM) possibly indicative of adaptation of the germline repertoire to this antigen. The presence of self-reactive B cell clones in GCs, formed in response to an immunodominant viral antigen, strongly supports a break of tolerance at the cellular level. Taken together, these findings indicate the potential relevance of NS1 for ZIKV pathogenicity and its associated autoimmune manifestations.

## Results

### Immunocompetent BALB/c mice develop a specific antibody response to ZIKV infection focused on NS1

To study the humoral immune response to ZIKV infection in immunocompetent mice, we injected young adult BALB/c WT mice intravenously with 10^7^-10^8^ PFU of the Brazilian ZIKV isolate PE243 [40] and followed antibody responses for 50 days post-infection (d.p.i) (Fig. 1 A). Mice showed increased spleen weight from day 7 to 28 after infection (Fig. 1 B), as well as altered total serum immunoglobulin concentrations. Increased serum IgM was detected at 7 d.p.i. (Fig. 1 C) and total serum IgG concentration increased progressively between 7 and 21 d.p.i., stabilizing at a higher concentration than controls for up to 50 days after infection (Fig. 1 D).

**Figure 1.**
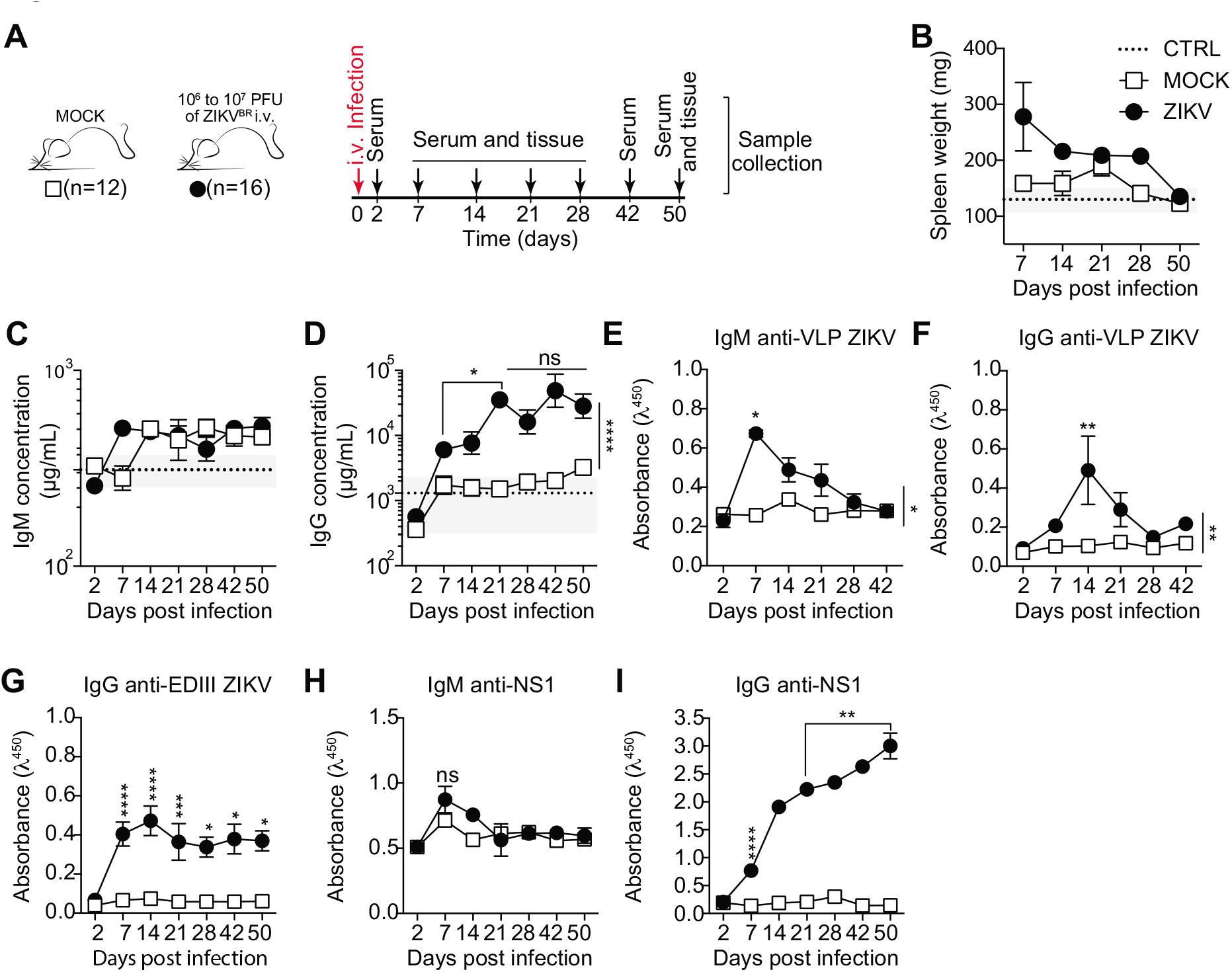
Characterization of ZIKV infection in immunocompetent BALB/c mice. **(A)** Experimental design indicating the time points of serum samples and lymphoid tissue collections. **(B)** Spleen weight measured at the time of collection, as indicated. Total serum IgM **(C)** and IgG **(D)** from infected (ZIKV) and control (MOCK) mice, measured by ELISA. (**E-G**) Levels of IgM and IgG specific for viral surface antigens were measured by ELISA (1:120 dilution) utilizing VLPs and recombinant domain III of ZIKV envelope protein (EDIII). (**H-I**) Levels of IgM (**H**) and IgG (**I**) specific for NS1 protein were measured by ELISA (1:120 dilution) utilizing recombinant ZIKV NS1. Data representative of three experiments with eight to sixteen mice per group. ns, not significant; *p<0,05; **p<0,01; ***p<0,001; ****p<0,0001.

Serum IgM binding to envelope proteins peaked at 7 d.p.i. (Fig. 1 E) followed by a peak in IgG at 14 d.p.i., (Fig. 1 F) as detected using Zika virus-like particles (VLPs) that display proteins E and M in their mature form [41]. Domain III of protein E (EDIII) is a target of neutralizing antibodies for different flaviviruses [34, 42-44]. We therefore also assayed serum for IgG binding to this portion of the E protein. We found that EDIII-specific IgG peaked in serum at 14 d.p.i. (Fig. 1 G), at which point serum neutralizing activity was also observed (data not shown).

We next searched for NS1-binding IgM and IgG antibodies in sera of infected mice. NS1 is known to be abundantly secreted into the extracellular milieu by flavivirus-infected cells [45] and is highly immunogenic [8]. IgM binding to NS1 remained almost unchanged compared to uninfected controls (Fig. 1 H). Interestingly, although serum IgG specific to NS1 appeared later than that targeting envelope proteins, anti-NS1 IgG increased progressively after infection, remaining at very high levels for as long as 50 d.p.i. (Fig. 1 I). Since viral RNA was undetectable in blood and brain tissue, in order to investigate whether our observations were dependent on viral replication leading to secretion of NS1 protein by infected cells, we performed the same experiment with UV-inactivated virus. Mice were injected intravenously with 10^7^-10^8^ PFU of UV-inactivated (iZIKV) or replicative ZIKV isolate PE243 [40] and evaluated for 60 days after infection (Fig. S1 A). UV-inactivated ZIKV did not induce an increase in spleen weight (Fig. S1 B). Despite the presence of E protein-specific IgG in serum (Fig. S1 C), these immunoglobulins did not target domain III (Fig. S1 D), and NS1-specific IgG was not detected in serum (Fig. S1 E). Taken together, these results demonstrate that, although immunocompetent mice survive ZIKV infection with few or no clinical signs, infection with replicative virus induced a robust humoral immune response, which became progressively dominated by antibodies to the NS1 antigen.

### Dominance of NS1-binding IgG in serum correlates with the emergence of autoreactive antibodies

The increasing levels of anti-NS1 IgG antibodies from 21 d.p.i. onwards prompted us to further investigate the NS1-specific response in infected mice. Serum titration at different time points suggested an increase in concentration or affinity of IgG for the viral antigen (Fig. 2 A). Endpoint titers increased gradually to approximately 200,000 at the latest time point assayed (Fig. 2 B). Serum IgG was predominantly of the IgG2a isotype throughout infection, as expected for anti-viral responses [46]. Of note, there was a late contribution of IgG1 to total serum IgG titer (Fig. 2 C and D). Given that gamma 1 constant region gene is located upstream of gamma 2a, ruling out sequential switching between these isotypes, our data suggest continued engagement of B cell clones after the initial phase of the response. We also assessed the binding of IgG to closely-related dengue virus antigens. As observed in human antibody response to ZIKV infection [15, 47], we found cross-reactivity between ZIKV EDIII and DENV EDIII (Fig. 2 E; Fig. S2, A and B), whereas ZIKV NS1-specific IgG did not cross-react with DENV NS1 protein (Fig. 2 F; Fig. S2, C and D).

**Figure 2.**
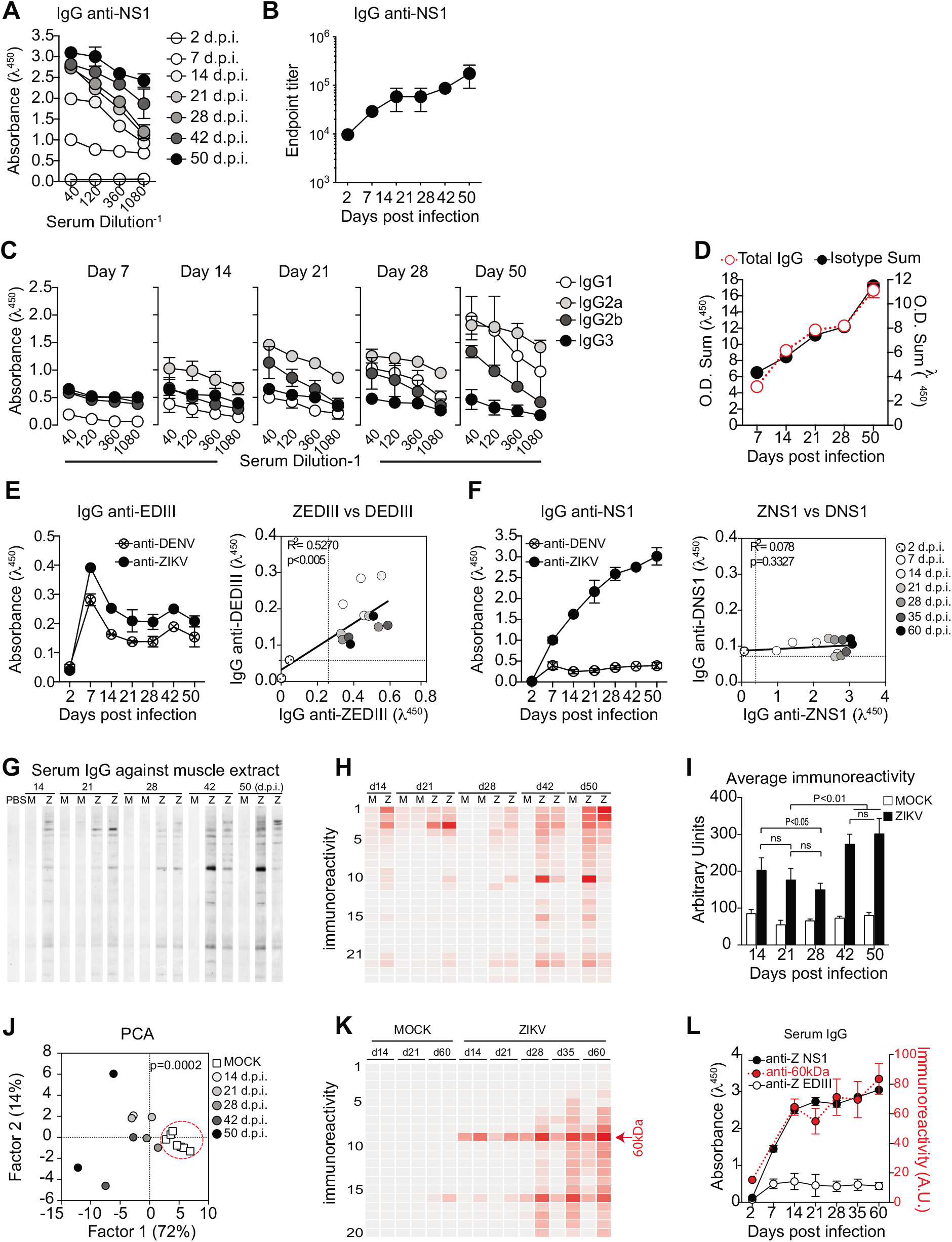
Humoral immune response to NS1 during ZIKV infection correlates with autoreactive antibodies. **(A)** Binding of serum IgG to ZIKV NS1 protein during infection detected by ELISA. **(B)** Endpoint titer of serum IgG specific to ZIKV NS1 protein during infection. **(C)** NS1-specific serum IgG isotypes composition during experimental infection were detected by antigen-specific ELISA. **(D)** Total NS1 specific IgG in serum corresponds to the sum of IgG isotypes, present in distinct proportions after infection. **(E-F)** Sera from ZIKV infected mice (1:120 dilution) were tested by ELISA for binding to ZIKV and DENV antigens EDIII (E) and NS1 (F). Correlation coefficients show cross-reactivity of EDIII–specific IgG but not of NS1-specific IgG. **(G)** Self-reactivities present in serum IgG (diluted 1:100) from control (M) and infected (Z) mice using muscle extract from Balb/c mice as source of self-antigens. **(H)** Intensity of bands was quantified and plotted as a heat map. **(I)** Total immunoreactivity (sum of all bands intensities) present in the sera of infected (ZIKV) and control (MOCK) mice for each time point after infection. **(J)** Principal component analysis (PCA) of all self-reactivities at all time points. Red circle indicates the segregation of the control group. **(K)** Intensity of reactivities present in serum IgG (diluted 1:100) from control (MOCK) and infected mice (ZIKV) using HEp-2 cell extract as source of self-antigens. **(L)** Intensity of the reactivity to a selected 60 kDa self-antigen throughout time after infection correlates with levels of serum IgG specific to ZIKV NS1 protein but not with levels of IgG specific to domain III of ZIKV envelope protein (ZEDIII). Data for A and B are from one experiment with three mice per group. Data for C and D are from one experiment representative of two experiments with five to sixteen mice per group. Data for E, F, K and L are from one experiment representative of two experiments with two to three mice per group. Data for G-J are from one experiment representative of two experiments with two representative mice per group. ns, not significant. ns, not significant.

In addition to the presence of virus-specific antibody responses, viral infections are often associated with hyperglobulinemia due to non-specific polyclonal activation of B lymphocytes [1]. To assess the potential of ZIKV infection to induce a polyclonal, non-specific humoral immune response, we looked for IgG binding to unrelated antigens such as heat shock proteins from both mammalian and commensal bacterial origins, which are commonly targeted by autoantibodies in different systems [48-50] (Fig. S2, E and F). IgG binding to unrelated antigens, possibly due to polyreactivity, was present at early time points and rapidly decayed, following the kinetics of antibody response to viral structural proteins shown in Figure 1. Interestingly, serum IgG from ZIKV-infected mice also displayed widespread binding to self-antigens, as revealed using a semiquantitative immunoblot assay that enables global analysis of the self-reactivity of antibodies present in serum [51, 52]. Accordingly, at 14 d.p.i., ZIKV-infected mice exhibited serum IgG reactivity to multiple self-antigens from syngeneic brain and muscle tissues (Fig. 2 G), suggesting a break in self-tolerance during the early humoral immune response.

Polyclonal B cell activation and non-specific polyreactive responses are mostly present in the acute phase of the immune response to viral infections, and do not contribute meaningfully to serum IgG titers at later time points when infection subsides. These observations prompted us to evaluate whether self-antigen reactivity declines at later time points after infection. Strikingly, serum IgG self-reactivity was progressively stronger at later time points (Fig. 2, G-I). Of note, although different mice shared several reactivities towards antigens with similar migration patterns (Fig. 2, G and H), individual immunoreactivity profiles did not necessarily converge in time towards a unique reactivity profile (Fig. 2 J).

To further investigate the progressive increase of self-reactive IgG following infection, we tested serum samples from an additional cohort of ZIKV-infected mice for binding to a HEp-2 cell line extract. These cells are frequently used in standard clinical assays for anti-nuclear antibodies (ANAs), but also as a source of cytoplasmic self-antigens [38]. Consistently, a broad range of immunoreactivities arose over time after infection (Fig. 2 K). The levels of IgG reactive to a 60 kDa self-antigen paralleled those of NS1-specific, but not of EDIII-specific, IgG over time (Fig. 2 L). The overlapping kinetics of anti-NS1 response and self-reactive serum IgG led us to hypothesize there could be a link between the maintenance of autoreactive antibodies and a dominant and sustained anti-NS1 antibody response.

### Virus-specific and autoreactive B cell clones are present in GCs after ZIKV infection

Viral infections typically induce T-dependent antibody responses, in which follicular B cells (FO) enter GC reactions where they undergo clonal expansion, SHM, and selection, leading to antibody affinity maturation (reviewed in [53]). After intravenous (i.v.) ZIKV infection, we observed abundant GC formation in spleen. Frequencies of different splenic B cell populations were found to be altered; the reduction in frequency of FO B cells likely reflected GC formation and correlated with levels of serum IgG specific to ZIKV proteins. GC B cell frequency peaked at day 14 after infection and started to decrease by day 21, receding to background levels at 28 d.p.i. (Fig. 3 A).

**Figure 3.**
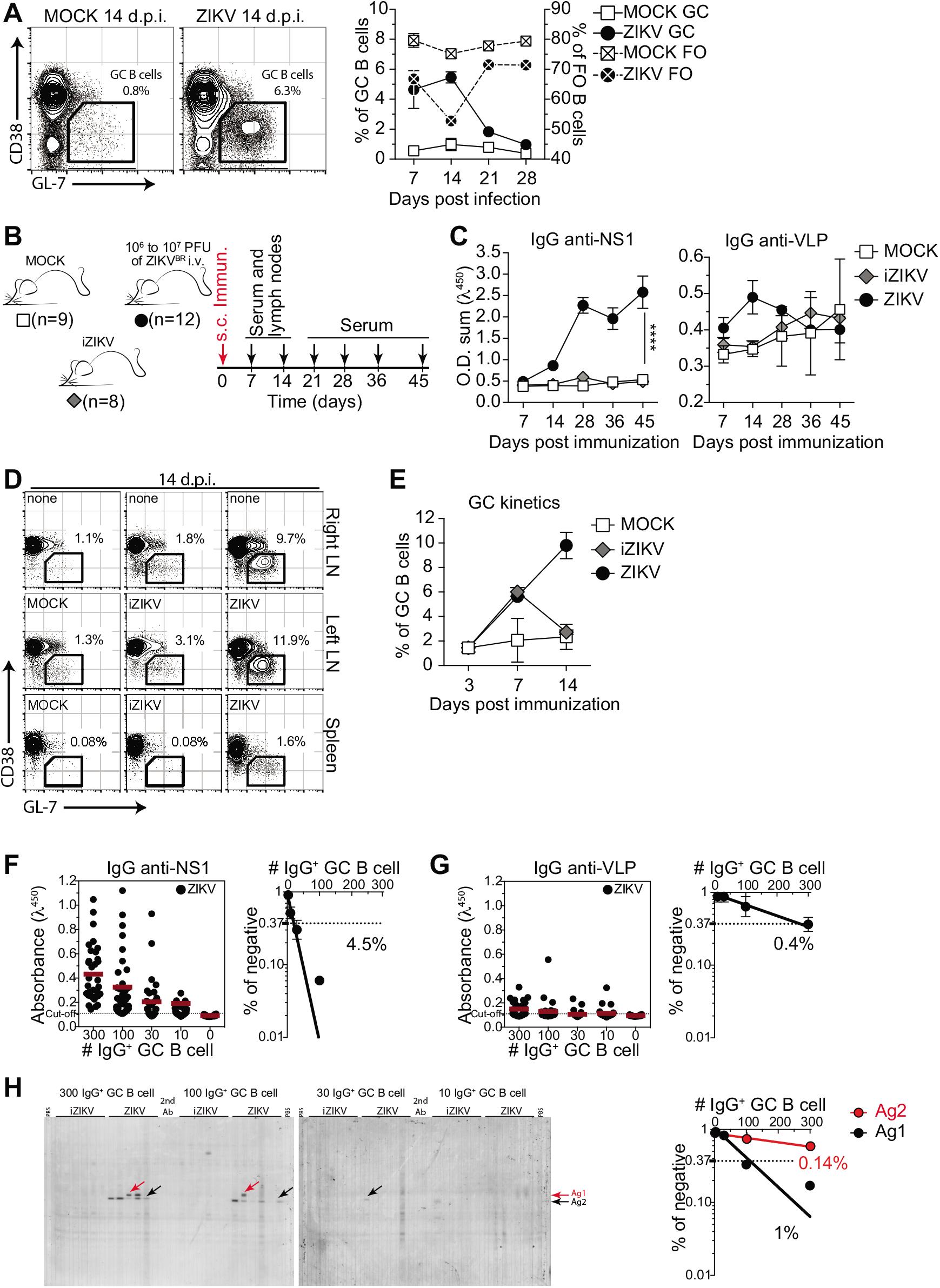
GC B cells produce both virus-specific and autoreactive antibodies. **(A)** GC B cells (CD38^lo/-^ GL-7^+^ gated on B220^+^ CD138^-^) in the spleen of ZIKV infected mice at 14 days post infection (left). Kinetics of frequency of follicular (FO) and germinal center (GC) B cells after infection is represented on right panel. (**B**) Subcutaneous infection experimental design indicating the time points of serum samples and lymphoid tissue collections from control mice (MOCK), mice immunized with UV-inactivated virus (iZIKV) and infected mice (ZIKV). (**C**) Kinetics of serum IgG specific to ZIKV NS1 and VLPs. OD Sum is the summation of ODs of four serum dilutions (1:40, 1:120, 1:360 and 1:1080). (**D**) Representative plots of GC B cells (CD38^lo/-^ GL-7^+^ gated on B220^+^ CD138^-^) at day 14 post infection. Mice were injected in the left footpad. (**E**) Kinetics of frequency of germinal center (GC) B cells in left popliteal lymph nodes after infection (ZIKV) or immunization (iZIKV). (**F-H**) GC B cells from popliteal lymph nodes of infected mice were sorted, pooled and cultured in decreasing numbers/well (300, 100, 30, 10 cells/well). Supernatants were collected on day 7 and screened for IgG secretion by ELISA. Supernatants that revealed the presence of IgG were tested for antigen specificity by ELISA **(F and G)** or immunoblot against mouse brain tissue as source of self-antigens **(H)**. Frequencies of IgG^+^ GC B cells that bound NS1 **(F)**, VLP **(G)** or self-antigens **(H)** were calculated utilizing Poisson distribution. Self-antigen reactivities used for frequency determination are indicated by arrows. Cell culture was performed on a monolayer of gamma-irradiated (20 Gy) NB40L feeder cells (3 × 10^3^ cells/well), LPS (30 µg/mL) and IL-21 (2 ng/mL). Data for A are from one experiment representative of two experiments with eight to sixteen mice per group. Data for D-E are from one experiment representative of two experiments with six to nine mice per group. Data for Fand G are from one experiment representative of two with three mice per group. Data for H are from a single experiment with three mice per group pooled for each sample.

To investigate the possible link between the antibody response to NS1 and the presence of self-reactive IgG in serum, we devised an experimental strategy that allowed us to more appropriately compare GC B cells between infected and immunized mice. For that purpose, mice were subcutaneously infected in the footpad with ZIKV and draining lymph nodes (LN) were collected on different days post-infection to isolate GC B cells (Fig. 3 B). Consistent with our previous results, we found increasing levels of IgG binding to ZIKV NS1 in serum up to 45 days after subcutaneous infection. This increase was not observed when the same amount of UV-inactivated virus (iZIKV) was injected, indicating that this phenomenon requires replicative ZIKV infection (Fig. 3 C). We could also observe GC formation in LN after both ZIKV infection and iZIKV immunization, although the latter were both of lower magnitude and shorter in duration (Fig. 3, D and E). At 14 d.p.i., corresponding to the peak of the response, we isolated and cultured GC B cells for Ig production *in vitro*. GC B cell cultures were performed as described by Kuraoka et al.[54], with modifications, including not adding IL-4 to prevent *in vitro* class switching. As a result, the proportions of IgG isotypes found in culture supernatants broadly matched those of ZIKV-specific antibodies in serum (Fig. S3).

Using limiting dilution analysis (LDA), we were able to estimate the frequency of GC B cells secreting immunoglobulins binding to VLP or NS1 (Fig. 3, F and G), as well as to self-antigens (Fig. 3 H). Quantification of the number of responding B cell clones per culture ensures accurate determination of the frequency of GC B cells reactive to a given antigen [55, 56]. GC B cells from mock-, iZIKV-, and ZIKV-injected mice showed similar frequencies of response to polyclonal LPS stimulus, with 30% to 50% of GC B cells proliferating and differentiating into IgG-secreting plasma cells in all conditions (Fig. S3 A, and data not shown). At 14 d.p.i., no reactivity to viral surface antigens was detected in GC B cells derived from control mice (mock) and less than 0.1% of GC B cells from iZIKV-imumunized mice secreted IgG that bound detectably to VLP. Moreover, we were unable to detect GC B cell clones secreting NS1-reactive IgG in neither mock-nor iZIKV-immunized mice (data not shown). On the other hand, in ZIKV-infected mice, GC B cells specific for NS1 were readily detected at relatively high frequencies (Fig. 3 F). The proportion of GC B cells reacting to NS1 (4.5%) was 10-fold higher than that of GC B cells binding to envelope proteins (0.4%) (Fig. 3 G) and correlated with virus-specific IgG levels observed in serum (Fig. 3 C; see also Fig. 1, F and J). These findings underscore the extent of the immunodominance of NS1 over envelope antigens also at the cellular level.

We then performed the global analysis of self-reactivities, as used for serum IgG (Fig. 2 H), with GC B cell culture supernatants. By combining LDA and immunoblot assays, we were able to estimate the frequency of cell secreting IgG binding to self-antigens in each GC B cell culture (Fig. 3 H). Our analysis showed that autoreactive B cells were present within GCs formed after exposure to replicative ZIKV, whereas these were virtually absent upon exposure to UV-inactivated virus (Fig. 3 H). Importantly, the self-reactivity found with highest frequency (Ag2) accounted for 1% of GC B cells in the lymph node (Fig. 3 H), at least twice the frequency of GC B cells detectably specific for virus envelope proteins (Fig. 3 G). These data suggest a role for NS1 in triggering the autoreactive antibody response following ZIKV infection.

### ZIKV NS1 immunization recruits a high frequency of NS1-specific B cells to the GC

To directly test whether an anti-NS1 humoral immune response generates autoreactive antibodies, we immunized mice in the footpads with purified recombinant ZIKV NS1 in the presence of a TLR7 agonist (R848). For comparison, two other groups were included. Mice were immunized with ZIKV virus-like particles (VLPs), displaying envelope proteins (E and M) in the presence of R848 or with a combination of VLPs and NS1 (Fig. 4 A). We first analyzed the humoral immune response against the viral antigens. We found that, from day 14 post-immunization onwards, serum levels of NS1-specific IgG were higher than those of VLP-specific IgG in both combinations (Fig. 4 B). Despite the similar frequencies of total GC B cells in popliteal LNs (Fig. 4 C), frequencies of specific B cells within the GC varied. At day 14 after immunization with VLP/R848, the frequency of GC B cells binding to VLPs was 2.6% (Fig. 5 E, left panels). After immunization with NS1/R848, however, frequency of specific GC B cells was approximately 10-fold higher, at 27% (Fig. 4 E, right panels). Compared to the single antigen immunization protocol, VLP and NS1 in combination reduced the frequency of GC B cells specific for both antigens to 1.2% and 3%, respectively. (Fig. 4 E, middle panels). This reduction was not due to lower secretion of IgG *in vitro*, since similar levels of NS1 specific IgG were found in culture supernatants, irrespective of the presence of VLPs in the immunization (Fig. 4 F). Altogether, antigen-specific GC B cell frequency and kinetics in immunized mice mirrored the virus-specific IgG levels found in the ZIKV infection model, corroborating the observed immunodominance of NS1 over ZIKV envelope antigens.

**Figure 4.**
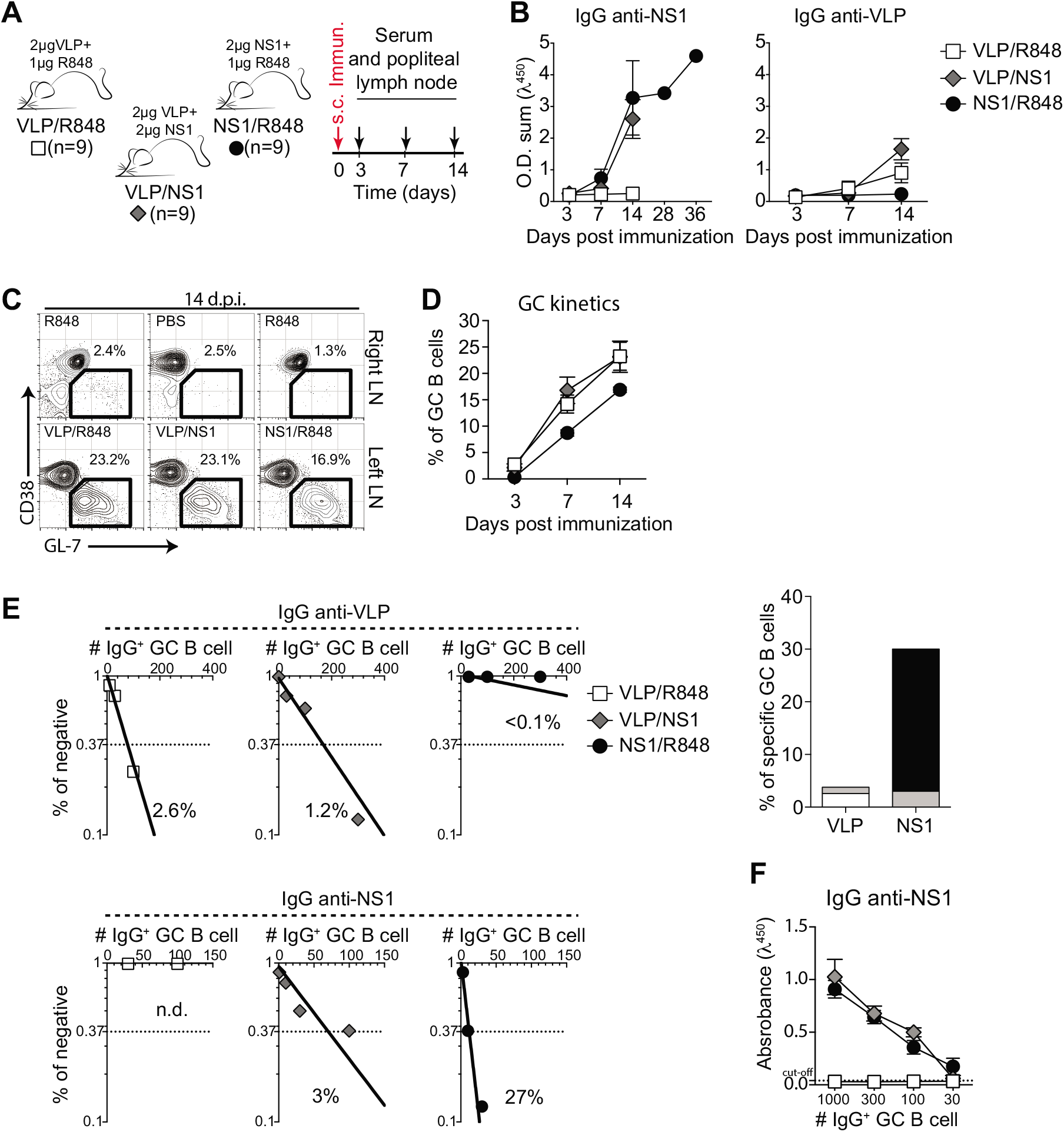
Antigen-specificity of B cells in germinal centers after immunization with ZIKV VLP and NS1. **(A)** Experimental design indicating the time points of serum samples and popliteal lymph nodes collections after immunization with NS1 (2 ug/mouse), VLP (2 ug/mouse) or both (2 ug of NS1 and 2 ug of VLP/mouse). Immunizations were adjuvanted with R848 (1ug/mouse). **(B)** Kinetics of serum levels of IgG binding to ZIKV NS1 recombinant protein or ZIKV VLP, measured by ELISA. OD Sum is the sum of ODs of four serum dilutions (1:40, 1:120, 1:360 and 1:1080). (**C**) Representative plots of GC B cells (CD38^lo/-^ GL-7^+^ gated on B220^+^ CD138^-^) at day 14 post immunization. Mice were immunized in the left footpad. (**D**) Kinetics of frequency of germinal center (GC) B cells in left popliteal lymph nodes after immunization. (**E**) GC B cells from popliteal lymph nodes of immunized mice were sorted and cultured in decreasing numbers/well (300, 100, 30, 10 cells/well). Supernatants were collected on day 7 and screened for IgG secretion by ELISA. Supernatants that revealed the presence of IgG were tested for antigen specificity by ELISA. Frequencies of IgG^+^ GC B cells that bound VLP (upper panel) or NS1 (lower panel) were calculated utilizing Poisson distribution and are summarized on the right graph. Cell culture was performed on a monolayer of gamma-irradiated (20 Gy) NB40L feeder cells (3 × 10^3^ cells/well), LPS (30 µg/mL) and IL-21 (2 ng/mL). (**F**) OD of IgG^+^ supernatants of different cell numbers/well binding to ZIKV NS1, measured by ELISA. Data representative of at least two independent experiments with nine mice per group.

### Paucity of somatic hypermutations in expanded GC B cell clones from ZIKV NS1-immunized mice

To further characterize the B cell response to ZIKV NS1, we sorted single GC B cells from popliteal LNs of mice immunized with this antigen at different time points and performed *Igh* sequencing. To gain insight into specific features of ZIKV NS1 B cell response, we also immunized mice with DENV NS1 for comparison, since, even though the two proteins are structurally homologous, antibodies generated after infection with ZIKV did not cross-react with DENV NS1 (see Fig. 2 F). As expected, GCs found in popliteal LNs of mice immunized with either ZIKV or DENV NS1 proteins 10 days post-immunization were highly clonally diverse, whereas clones with increased frequency accumulated over time (Fig. 5A).

**Figure 5.**
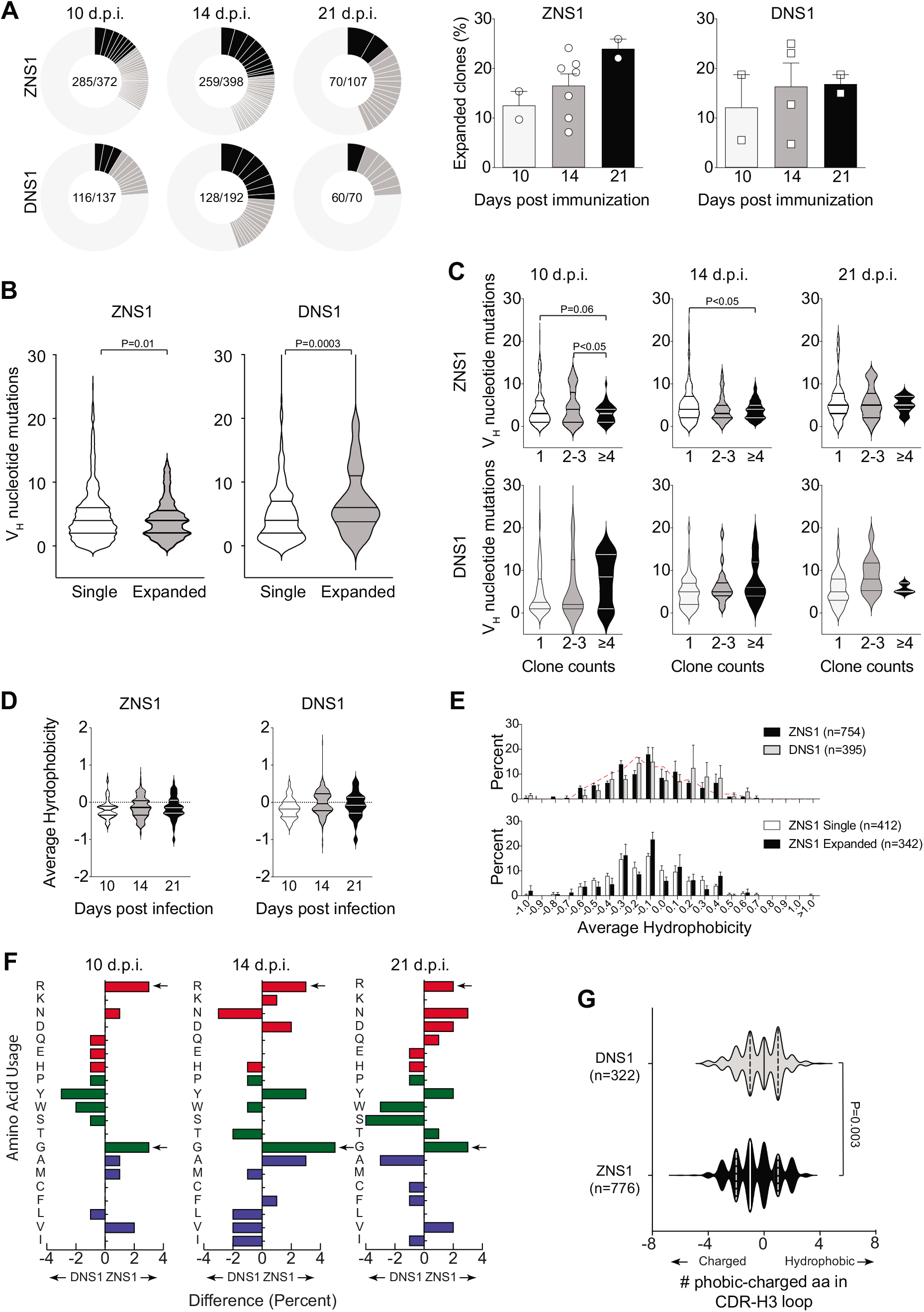
Characterization of B cell repertoire present in GCs after ZIKV NS1 immunization. **(A)** Single GC B cells from mice immunized s.c. with recombinant ZIKV NS1 or DENV NS1 were sorted at different time points after immunization and *Igh* gene was sequenced. Pie charts represent clonal diversity found in all lymph nodes analyzed. Slices represent clonotypes assigned based on V_H_ and J_H_ usage and CDRH3 length and sequence. Slice size is proportional to the frequency of each clone. Black slices indicate clone counts higher than 4. Dark gray slices indicate clone counts of 2-3. Light gray where slices are not delimited represents single clones. Proportion of expanded clones is indicated on the right. **(B)** Number of somatic mutations found in V_H_ segments separated by singletons vs expanded clones. **(C)** Number of somatic mutations found in V_H_ segments at different time points separated by clone count. **(D)** CDR-H3 average hydrophobicity index variation among all sequences at different time points after immunization. (**E**) Comparison of CDR-H3 average hydrophobicity index distribution among all sequences from mice immunized with ZIKV NS1 or DENV NS1 (upper panel) and comparison between expanded (clonotypes found more than once in the same lymph node) and single clones from mice immunized with ZIKV NS1 (lower panel). Dashed red line indicates the distribution of CDR-H3 average hydrophobicity in follicular B cells from WT BALB/c mice. The normalized Kyte–Doolittle hydrophobicity scale [89] was used to calculate average hydrophobicity. (**F**) Divergence in the distribution of individual amino acid use in the CDR-H3 loop between ZNS1 versus DNS1 immunized mice at different time points. Red bars indicate charged amino acids, green bars represent neutral amino acids and blue bars, hydrophobic amino acids. (**G**) Difference in number of hydrophobic and charged amino acids was calculated for each CDR-H3 sequence and distribution was plotted for all sequences from DENV NS1 immunized mice (gray) and ZIKV NS1 immunized mice (black). Lymph nodes were sorted and sequenced individually from 2-5 mice per group at each time point. Sequences were pooled for analyses.

Analysis of V_H_ segment usage showed preferential use of the V_H_1 (J558) gene family, irrespective of immunizing antigen and time point analyzed (Fig. S4 A). The average number of mutations in V_H_ gene segments was similar for both antigens and increased over time as one would expect. However, interestingly, V_H_ mutation numbers were lower among clones found more frequently (likely those undergoing positive selection) in GCs formed after ZIKV NS1 immunization when compared to those formed after DENV NS1 immunization (Fig. 5 B). In contrast to those clones from ZIKV NS1 immunization, B cells in GCs from DENV NS1-immunized mice tended to progressively accumulate mutations in more expanded clones over time (Fig. 5 C).

Although no significant differences in CDR-H3 length were found in GC B cells after DENV or ZIKV NS1 immunization at any time point (Fig. S4 B), there was a preference for 11-amino acid-long CDR-H3s in both ZIKV NS1 and DENV NS1 immunized mice when compared to the naïve B cell repertoire (Fig. S4 C). Average CDR-H3 hydrophobicity tended to be lower for ZIKV NS1 than for DENV NS1 (Fig. 5 D). The distribution of CDR-H3 average hydrophobicity revealed few highly charged sequences among B cells responding to ZIKV NS1 antigen (Fig. 5 E). Comparison of the amino acid composition of CDR-H3 regions of clones obtained in response to both antigens at different time points revealed an enrichment over time for charged amino acids, especially arginine after ZIKV NS1 immunization (Fig. 5 F). Of note, charged amino acids in the antigen binding sites of IgG are often critical for self-reactivity [57]. However, a marked glycine (neutral amino acid) enrichment was also observed, which could counterbalance the charged amino acid bias, resulting in a moderate change in average hydrophobicity in CDR-H3 sequences as shown in Fig. 5 D. Evaluating the percent difference between hydrophobic and charged amino acids in CDR-H3 sequences, we found that B cell clones derived from the ZIVK NS1 immunized animals used charged amino acids more frequently than hydrophobic ones when compared to the DENV NS1-immunized mice (Fig. 5 G).

The recruitment of B cells enriched in charged CDR-H3 sequences into the GCs upon immunization with ZIKV NS1 could be related to the emergence of autoreactive IgG. Although, the specificities of the immunoglobulins encoded by these sequences are not known, results obtained with GC B cells cultures suggest that most of these cells (63% for ZIKV NS1 and 97% for ZIKV VLP) do not secrete antibodies with detectable binding to the antigen used in immunization (see Fig. 4 E). One might expect that the most expanded clones would be those with the greatest affinity for the antigen; however, it remains possible that the most mutated clones would become expanded only at later time points, after more extensive selection, and would therefore not be detected at high frequencies in the samples we analyzed. To better understand this phenomenon, we sought to correlate the features of immunoglobulin variable gene sequences with the specificities of their secreted antibodies.

### ZIKV NS1 immunization recruits into GCs B cell clones enriched for charged amino acids in the CDR-H3s and self-reactivity

To understand the relationship between Ig sequence features and antigen specificity, we used single GC B cell cultures, which allow the assessment of both *Ig* sequence and binding properties from the same cell (adapted from [56]). For this purpose, GC B cells were harvested from popliteal LNs of mice immunized with ZIKV or DENV-NS1 protein at different time points after immunization. After 7 days in culture, cells were processed for *Igh* sequencing and supernatants were used for specificity assessment. We first determined the frequency of GC B cells secreting ZIKV NS1-binding IgG. A total of 271 monoclonal antibodies were tested for binding to NS1 protein. In line with the rising levels of serum IgG specific for NS1 (shown in Fig. 4 B), frequency of ZIKV NS1-specific B cells also increased over time, from an average of 30% at 10 d.p.i. to 50% at 21 d.p.i. (Fig. 6 A). We then divided B cell clones into NS1-binders and non-binders for further analyses. Clonal expansion was evident among NS1-binding GC B cells, peaking on day 14 after immunization (when almost 60% of NS1-binders were found in detectably expanded clonotypes), consistent with antigen-driven selection (Fig. 6 B). At 21 d.p.i., the fraction of expanded clones among NS1-binders decayed to a frequency similar to that found at day 10 post-immunization. Although the number of clones analyzed at this time point is limited, this result could suggest continued ingress of new B cell clones into ongoing GCs (Fig. 6 B). In contrast, non-binder clones showed significantly less clonal expansion at all time points analyzed (Fig. 6 B). Interestingly, and consistent with data shown previously (Fig. 5 B), we did not find evidence for positive selection of NS1-binding clones bearing large numbers of somatic mutations (Fig. 6 C).

**Figure 6.**
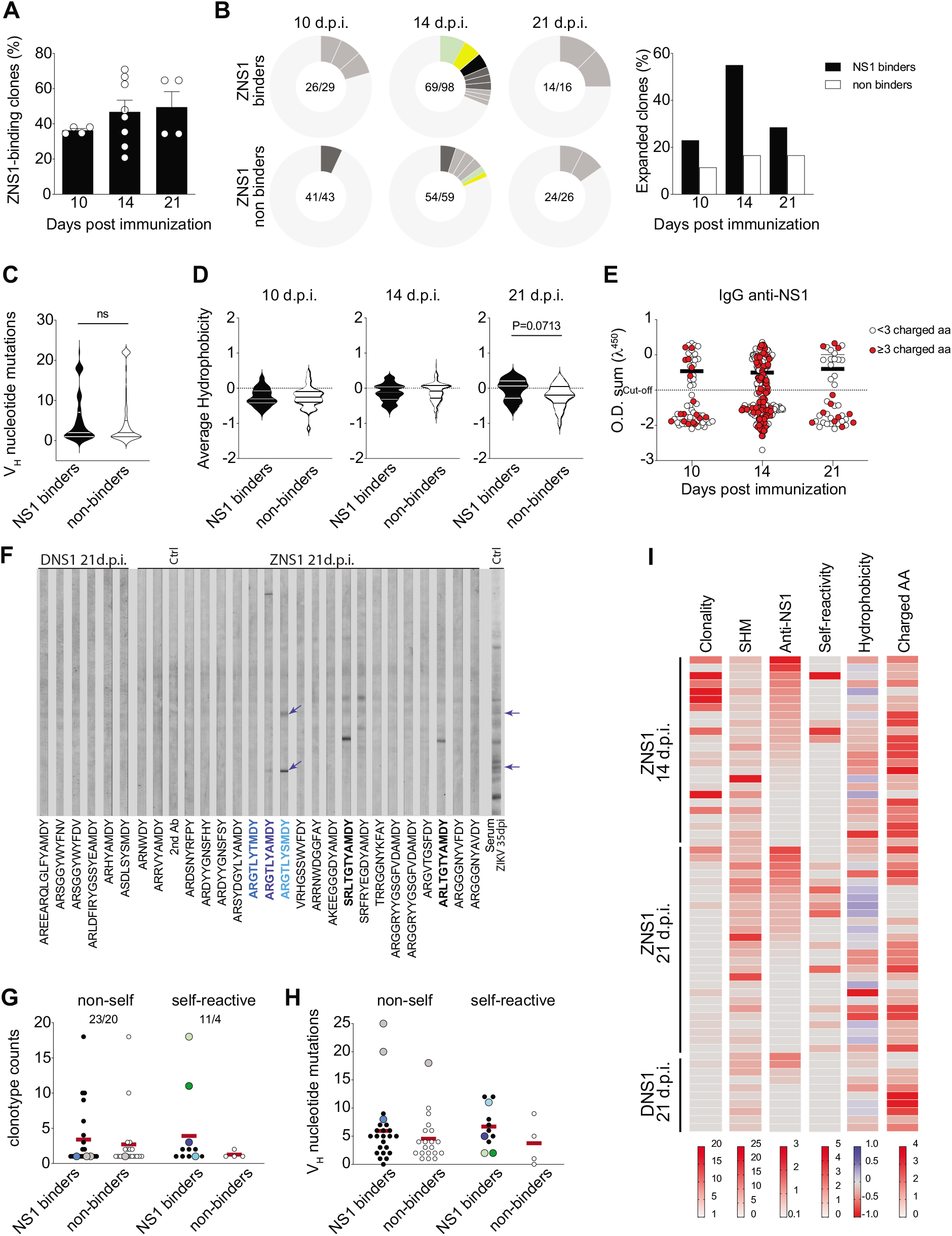
Self-reactivity of GC B cells after ZIKV NS1 immunization. Single GC B cells were sorted and cultured on a monolayer of gamma-irradiated feeder cells (1×10^3^ cells/well) expressing CD40L, BAFF and IL-21 (Kuraoka et al., 2016). After 7 days, supernatants were collected for binding assays and cells were harvested for *Igh* sequencing. **(A)** Frequency of IgG^+^ single GC B cell culture supernatants that bound to ZIKV NS1. Supernatants were screened for IgG production and IgG^+^ wells were tested for binding to ZIKV NS1 protein by ELISA. **(B)** Clonal distribution of GC B cells found to bind to NS1 (upper panel) or that did not bind to NS1 (lower panel). Size of the slice is proportional to the clone frequency. Colored slices represent variants of clones that were found both as binders and non-binders. Right panel represents the frequency of expanded clones among binders and non-binders at specific time points after immunization. **(C)** Number of somatic mutations found in V_H_ segments from each GC B cell sequenced grouped based on binding to NS1. **(D)** CDR-H3 average hydrophobicity index variation at different time points grouped by binding to NS1. **(E)** NS1 binding by ELISA OD related to the presence of charged amino acids. Red dots indicate the presence of 3 or more charged amino acids in CDR-H3 at different time points. **(F-I)** Single GC B cell culture supernatants were tested for binding to self-antigens by immunoblot. **(F)** Representative immunoblot profiles of monoclonal IgG from single GC B cell culture supernatants binding to mouse brain extract. Blue arrows indicate immunoreactivities highlighted in the main text **(G)** Clonotype counts of NS1 binders (black dots) and NS1 non binders (white dots) separated by self-reactivity. Color coded dots represent variants of clones described in the main text. **(H)** Number of somatic mutations found in V_H_ segments grouped by binding to NS1 and self-reactivity. Color coded dots represent variants of clones described in the main text. **(I)** Clonality, somatic hypermutation, binding to NS1, self-reactivity, hydrophobicity and charged amino acid usage by time after immunization. Clonality corresponds to the number of variants of each clonotype found in the dataset. SHM is represented by the number of V_H_ mutations found in each sequence. Anti-ZNS1 indicates the OD obtained by ELISA. Self-reactivity corresponds to the number of bands found for each supernatant in mouse tissue extracts (brain and/or muscle). Hydrophobicity corresponds to the average hydrophobicity of the CDR-H3 loop, hydrophobic and charged CDR-H3 sequences are shown in blue and red, respectively. Charged AA indicates the number of charged amino acids found in CDR-H3 loop. Lymph nodes were sorted and cultured individually from 2-3 mice per group at each time point. Sequences were pooled for analyses.

We then investigated the physicochemical features of the CDR-H3 sequences expressed by B cells recruited to GCs after ZIKV NS1 immunization. On day 14 p.i., both ZIKV NS1 binders and non-binders showed similar CDR-H3 hydrophobicity, whereas on day 21 p.i., ZIKV NS1 binders tended to be less charged and closer to neutrality than non-binders, an effect just short of statistical significance (Fig. 6 D). To examine whether NS1 binding correlated with the abundance of positively charged amino acids in CDR-H3, we plotted the anti-NS1 reactivity per clone at each time point, highlighting CDR-H3s expressing three or more charged amino acids (Fig. 6 E, red dots). CDR-H3 sequences bearing charged amino acids were frequent and evenly distributed among NS1-binder and non-binder clones, accounting for 25% to 35% of CDR-H3 sequences (Fig. 6 E). Overall, CDR-H3 sequences of GC B cells from ZIKV NS1 immunized mice were enriched for charged amino acids (including arginine), as compared to what has been observed in the naïve repertoire [58] as well as in comparison to clones from DENV NS1 immunized mice (Fig. 5 F and G; Fig. 6 E).

The presence of positively charged amino acids within CDR-H3 is a common feature associated with self-reactivity [59]. To determine whether particular *Igh* sequence features were correlated with self-reactivity, we tested 58 monoclonal antibodies for binding to autologous muscle and brain extracts (Table S1 and Fig. 6 F). Corroborating the results obtained with serum and limiting dilution of GC B cells, no self-reactivity was found among GC B cells from mice immunized with DENV NS1 at any time point (Table S1 and Fig. 6 F). In contrast, 18.2% of evaluated single cells obtained at day 14 after immunization with ZIKV NS1 were self-reactive. This frequency increased to 42.3% of the analyzed cells from 21 d.p.i. Highly charged CDR-H3 were not determinant for self-reactivity since we found this feature among both self-reactive and non-self-reactive clones. Nevertheless, we did find a handful of clonotypes bearing hydrophobic amino acids in their CDR-H3 that were predominantly self-reactive (Table S1; Fig. 6, F and I).

### GC B cells from ZNS1 immunized mice display widespread self-reactivity irrespective of clonal sizes, SHM and nominal antigen-selection

We further analyzed the SHM patterns and clonal frequencies of 58 GC B cells for which we obtained complete data set, correlating it with IgG reactivity to NS1 and to self-antigens. Among GC B cells obtained at day 14 after immunization with ZIKV NS1, we found cells with similar CDR-H3 but different SHM patterns, including a clonal family bearing a “glycine-enriched CDR-H3” (ARGGGYDGFAY). The least mutated cell in this group was inferred to have acquired a non-silent mutation in the D2-02 gene segment, which led to the replacement of a tyrosine by a phenylalanine in the CDR-H3 (becoming ARGGGFDGFAY). The germline D2-02 D gene segment encodes the GYD motif, whereas the mutated version encoded a GFD motif, which displayed increased poly-reactivity against self-antigens together with increased reactivity to ZNS1 (Fig. S5 A and Table S1). The two clones bearing the glycine-enriched CDR-H3 that bound NS1 and cross-reacted with self-antigens were also highly expanded, suggesting that antigen-specific GC B cells – whether self-reactive or not – are capable of undergoing positive selection (Fig. 6 G, green dots).

At 21 d.p.i., other groups of related clonotypes were found with different SHM patterns (Table S1 and Fig. 6 F). An interesting autoreactive clone carried the ARGTLYAMDY CDR-H3 (Fig. 6 F, highlighted in blue colors). In addition to being among the most hydrophobic sequences in our list, we found more mutated variants both non-autoreactive (ARGTLYTMDY) and autoreactive (ARGTLYSMDY) (Fig. S5 B, Fig. 6 F and Table S1). Although all variants of this clonal family bound to ZIKV NS1 protein, the most mutated one (ARGTLYSMDY) displayed the highest relative reactivity to NS1 (Table S1) as well as increased poly- and self-reactivity (Fig. S5 B and Fig. 6 F, highlighted in light blue). In this case, only the least mutated sequence (ARGTLYAMDY) was found more than once in the data set. Notably, the two most prominent self-reactivities found in the ARGTLYSMDY clone were also found in serum IgG from mice infected with ZIKV 35 d.p.i. (Fig. S5 B and Fig. 6 F, blue arrows).

Overall, ZIKV NS1 clonotypes found at the highest frequencies (> 5 counts) were generally non-self-reactive (Fig. 6 G). Moreover, all NS1-binding immunoglobulins tested for self-reactivity, regardless of their self-reactivity status, had numbers of V_H_ mutations that were similar and higher than those of non-antigen-specific cells, again suggesting antigen-driven selection (Fig. 6 H). The most mutated clones did not bind to self-antigens and were found exclusively as singletons, irrespective of binding to ZIKV NS1 (Fig. 6 G and H, gray dots).

The results obtained with single cell sequencing and binding assays were summarized in Figure 6 I. Clonality scores showed that the presence of more expanded clones was a property of earlier GCs, obtained at day 14 post-infection, whereas late GCs were enriched in smaller clones. Interestingly, the decrease in clonal dominance inversely correlated with increased self-reactivity, suggesting a replacement of early clones by a new wave of B cells with self-reactive potential. SHM appeared not to increase from day 14 to 21, which would be in line with clonal replacement. Self-reactive clones were already present at day 14, all of which exhibited cross-reactivity with ZIKV NS1. It is noteworthy that autoreactive clones with no cross-reactivity with NS1 were present at day 21 but not at day 14 p.i.. Although the number of clones assayed for autoreactivity limits the strength of this conclusion, we found more self-reactivity among NS1-binders than among non-binders. Collectively, our data support a role for NS1 in recruiting cross-reactive B cell clones into GCs, some of which are self-reactive clonotypes showing evidence for clonal expansion and paucity of VH gene mutations.

## Discussion

The humoral immune response to ZIKV infection in humans is characterized by early appearance of antibodies to structural proteins of the viral envelope followed by a later increase in antibodies to the non-structural protein NS1 [14, 15]. Here, we found that the humoral immune response of immunocompetent BALB/c mice to ZIKV infection follows a similar pattern. The BALB/c antibody response is characterized by early emergence of envelope-specific IgG, including EDIII-specific and neutralizing antibodies, followed by a delayed response dominated by antibodies to NS1. The delayed response to NS1 may be a consequence of its absence from the viral particle, possibly explaining the initial dominance of anti-envelope antibodies. Only after productive ZIKV infection do cells start producing and secreting large amounts of NS1, which then accumulates in bodily fluids and on the surface of infected cells. NS1 circulates in blood, and has been implicated both in damage to endothelial cells and in immune evasion through inhibition of the complement cascade [12, 23, 60, 61]. Of note, antibodies to NS1 have been shown to be protective against Zika and Dengue diseases, and immunization with NS1 has been considered as a possible prophylactic strategy [18, 19, 62].

Antibodies to DENV NS1 have also been implicated in pathological autoreactivity in humans [20, 63]. From this perspective, it is interesting to note that reactivity towards self-antigens present in different tissues or cell extracts was observed in the humoral immune response of BALB/c mice to ZIKV infection. The presence of autoreactive antibodies in viral diseases is not an uncommon finding and is often attributed to non-specific polyclonal B cell activation. Autoreactive B cells could be stimulated in a T-independent manner, via simultaneous TLRs and BCR signaling both in GCs and in extrafollicular foci [64-67]. However, polyclonal B lymphocyte activation is mostly an acute phase phenomenon that subsides with clearance of the virus. Here, by contrast, we found that self-reactivity was sustained and even augmented at later time points after infection, long after ZIKV had been eliminated. Interestingly, NS1-specific antibodies were also long-lasting in serum and correlated well with the kinetics of appearance and maintenance of autoreactive antibodies in BALB/c mice. Serum IgG isotypes composition of the NS1 specific repertoire was coherent with the sustained IgG anti-NS1 response and with engagement of new clonotypes. Early predominant IgG2a response was followed by a later emergence of IgG1, while IgG2a titers still increased up to 50 days after infection. As previously observed, gamma 1 constant region gene is located upstream of gamma 2a, ruling out sequential switching between these isotypes, implying the recruitment of new clones to sustain anti-NS1 antibody levels. It is still not clear whether the response to the viral antigen is maintained by self-antigens or by the viral antigen itself, that could be present at later time points, even though the virus was not detected after the first week of infection, as long-term antigen persistence in follicular dendritic cells has been well documented [68]. Hence, kinetics of the NS1 response is unique and differs from that induced by viral surface antigens, progressively dominating the humoral response. The shift from anti-envelope to anti-NS1 response could be a viral escape strategy, since neutralization is achieved mostly by antibodies directed to viral surface proteins.

Our analysis of GC B cells from ZIKV infected mice revealed both virus-specific and self-reactive B cells. Using single-cell cultures of GC B cells from mice immunized with ZIKV NS1, we showed that both anti-NS1 and autoreactivity could often be attributed to the same cell. Although GC responses to ZIKV NS1 and DENV NS1 did not differ in kinetics and were of short duration, autoreactive clones were found at all time points investigated after ZIKV NS1 immunization but not after DENV NS1 immunization. It is worth noting that serum IgG anti-ZIKV NS1 did not cross react with DENV NS1, in agreement with the distinct reactivity profiles reported here.

Self-reactive antibodies associated with autoimmune diseases, such as lupus, have been shown to be enriched for charged amino acids, especially arginine, in CDR-H3 [39]. Under normal circumstances, immunocompetent BALB/c mice are considered resistant to production of these autoantibodies [69, 70]. This is in part due to the low prevalence (around 5%) of arginine in the CDR-H3 region of the BCRs of mature recirculating B cells in BALB/c mice [58]. Half of the arginines in CDR-H3 regions of immature B cells in BALB/c mice derive from N-additions and half from germline D_H_ sequences [70]. Here, we found an enrichment for charged amino acids, including arginine, in CDR-H3 of GC B cell receptors after ZIKV NS1 immunization, as compared to DENV NS1. Interestingly, one particular clone found at high frequency among GC B cells at day 14 after immunization with ZIKV NS1 utilized the D gene segment DSP2.11 (D2-14), the only one to encode arginine in the germline sequence in reading frame 1 [70]. Notably, a marked glycine enrichment in CDR-H3 was also observed, which might contribute to maintaining the average hydrophobicity close to neutrality. NS1 is a complex multifunctional protein that forms a peculiar hydrophobic core in its hexameric structure which is secreted by infected cells [71]. Interestingly, in addition to the enrichment for charged amino acids, we also found self-reactive clonotypes bearing hydrophobic CDR-H3s; whether these particular clonotypes are able to bind to hydrophobic epitopes on NS1 protein, revealing potential immunogenicity of this hydrophobic core remains to be determined.

*Igh* sequencing of B cells isolated from GCs of ZIKV NS1 or DENV NS1-immunized mice showed similar usage of V_H_ segments and similar average number of V_H_ mutations, increasing over time as one would expect. However, after ZIKV NS1 immunization, V_H_ mutation numbers were lower among clones found more frequently, possibly indicating selection of near-germline B cell clones. We speculate that this could be due to the availability, in the pre-immune repertoire, of B cells able to bind to ZIKV NS1 protein with enough affinity to differentiate rapidly into plasma cells before accumulating many mutations. As recently shown by Burnett and colleagues, selection in GCs is skewed towards lower affinity for self-antigens prior to increasing affinity to foreign antigens [72]. In this context, it is possible that anti-ZIKV NS1 clones could generate plasma cells before acquiring enough mutations to diminish affinity to self-antigens. While it is known that plasma cell differentiation in GCs is dependent on high BCR affinity, the mechanisms through which the affinity threshold for differentiation is set are unclear [73, 74].

Coupling sequence analysis to binding assays, we found NS1-specific B cells that had similar numbers of V_H_ mutations (higher than those of non-specific clones), regardless of being self-reactive or not. By contrast, self-reactive B cells that did not bind to ZIKV NS1 protein were the least mutated of all and were also not detectably expanded (Figure 6 G and H). These data suggested the presence of antigen-driven selection in spite of self-reactivity, although, among NS1-specific cells, self-reactive cells seemed disfavored as compared to non-self-reactive ones. For instance, the glycine-enriched CDR-H3 ARGGGYDGFAY likely further mutated to ARGGG**<u>F</u>**DGFAY, generating increased poly- and self-reactivity while simultaneously increasing its reactivity to ZNS1. The autoreactive clonotype ARGTLY**<u>A</u>**MDY, on the other hand, was found in further mutated versions, ARGTLY**<u>T</u>**MDY, which lost autoreactivity, and ARGTLY**<u>S</u>**MDY, which remained autoreactive (Fig. S5 A and Table S1). Overall, our observations suggest that GC B cell clones undergoing selection after ZIKV NS1 immunization tend to be closer to germline than those in DENV NS1 immunization, an intriguing finding that requires further investigation.

In conclusion, we show here, for the first time, that ZIKV NS1-specific GC B cells can cross-react with self-antigens, possibly by molecular mimicry between ZIKV NS1 and self-antigens, raising the question whether self-antigens can participate in the stimulation of anti-NS1 B cell clones. This hypothesis could explain the sustained progression of the anti-NS1 humoral immune response we observed in infected mice, which display similar kinetics to that of self-reactive antibodies in serum. Notably, autoreactive clones that did not react with ZIKV NS1 were also found in GCs, raising the possibility of a “true” break of self-tolerance in the immune response to viral infection that goes beyond antigen mimicry. Finally, the flavivirus NS1 protein has been proven highly immunogenic in humans and capable of inducing protective antibodies and therefore suggested as a potential vaccine antigen [16, 18] or therapeutic antibody target [62]. In this context, the data presented here raise concerns about the safety of that approach and call attention to in-depth analysis of B cell clones engaged in response to this viral antigen, especially in its autoreactive component. This concern is echoed in very recent studies highlighting the self-reactive potential of close-to-germline encoded anti-viral antibodies to severe acute respiratory syndrome coronavirus 2 (SARS-CoV-2), the infectious agent of coronavirus disease 2019 (COVID-19) [75-78], arguing that the phenomenon we describe may be of importance well beyond the specific case of ZIVK infection.

## Material and methods

### Mice and treatments

BALB/c adult female mice, aged 6 to 8 weeks, were obtained from NAL-UFF, LAT-UFRJ or The Jackson Laboratory. Mice were kept in a 12h light/dark cycle with ad libitum access to food and water. Mice were infected with 10^6^-10^7^ PFU of ZIKV PE243 [40] i.v. and blood samples were collected on days 2, 7, 14, 21, 28, 35, 42, 50 or 60, as indicated for each experiment. ZIKV strain PE243 (Brazil/South America, gene bank accession no. KX197192) was propagated and titrated in Vero cells and UV-inactivated as previously described [40]. All animal procedures were approved by the Institutional Animal Care and Use Committee of the Centro de Ciências da Saúde da Universidade Federal do Rio de Janeiro and the Rockefeller University.

### ELISA for immunoglobulin quantification

Total IgG and IgM concentration in serum and cell culture supernatants were determined by ELISA as previously described in [79] using anti-mouse IgM and IgG-specific reagents (Southern Biotechnology). Briefly, 96 well plates (Costar) were incubated with anti-IgM or anti-IgG capturing antibody at 1ug/ml (Southern Biotech) and incubated at 4°C for 18h. Then, after 1h of blocking (PBS-1%BSA), serum samples and culture supernatants were diluted in PBS-BSA 1% by serial dilution starting at 1:40 for serum and undiluted for supernatants. Standard curves of polyclonal IgM or IgG were obtained by serial dilution 3-fold for IgM and 5-fold for IgG, starting at 1ug/mL for supernatants and 2ug/mL for serum samples. Secondary antibodies conjugated to HRP (Southern Biotech) were diluted 1:2000 for IgM and 1:8000 for IgG in PBS-BSA 1%. After wash with PBS, reactions were developed with TMB substrate solution (Sigma). The reaction was stopped with HCL 1N.

### ELISA for antigen-specific immunoglobulin detection

ELISAs to determine serum levels of anti-VLPs, anti-EDIII and anti-NS1 antibodies as well as specificity of IgG in B cell culture supernatants were performed as previously described[35]. Briefly, 96 well plates (Costar) were coated with peptide (EDIII) (10 µg/ml) or protein (1µg/ml) diluted in PBS and incubated at 4°C for 18h. Serum samples were diluted following serial dilution 1:3 for IgM and IgG starting with 1:40 in the first well. ZIKV and DENV EDIII recombinant proteins were kindly provided by Dr. Orlando Ferreira, IB, UFRJ. VLPs were produced and purified as previously described [41]. ZIKV and DENV NS1 proteins were produced and purified as reported previously.

### Immunoblot

BALB/c tissues (brain and skeletal muscle) were dissociated by Polytron homogenizer (4000 rpm) in homogenizing buffer (Tris-HCl 0,5M pH 6,8, SDS 10%, Mili-Q water) as described by Haury et al.[52]. Tissue extracts were fractioned by electrophoresis in 10% polyacrylamide gel under denaturing conditions, at 50mA until 6cm of migration. Proteins were transferred from the gel to a nitrocellulose membrane by a semi-dry electro transfer (Semi-Dry Electroblotter B) for 60 minutes at 0,8mA/cm^2^. After transfer, the membrane was kept in 50mL of PBS/Tween 20 (BioRad) at 0,2% vol/vol shaking for 18 hours at room temperature.

Incubation of the membrane with cell culture supernatants or serum samples was performed in a Miniblot System Cassette (Immunetics Inc.) which allows the simultaneous incubation of 28 different samples in separated channels. Supernatant samples were diluted 1:2 and serum samples were diluted 1:100. After wash, membranes were incubated with secondary antibodies conjugated to alkaline phosphatase anti-IgM or anti-IgG diluted 1:2000 (rabbit anti-mouse IgM, Jackson ImmunoResearch, goat anti-mouse IgG, SouthernBiotech). Substrate NBT/BCIP (Promega) was added after wash. Reaction was developed shaking at room temperature and stopped with Milli-Q water. Colloidal gold staining was performed after scanning membranes.

### Flow cytometry and cell sorting

Cells were harvested from spleen, peritoneal cavity and popliteal lymph nodes for flow cytometry and cell sorting. Splenocytes were homogenized with complete RPMI 1640 medium (GIBCO), followed by red blood cell lysis in 1 mL of ACK lysing buffer (GIBCO) for 1 min on ice. Single cell suspensions were prepared from the peritoneal cavity lavage with 5ml of complete RPMI 1640 medium (GIBCO). Popliteal lymph nodes were homogenized with complete RPMI 1640 medium (GIBCO). Cells were washed and resuspended in an appropriate volume for counting and staining. Cells were stained with the following monoclonal Abs conjugated to fluorochromes: anti-B220, CD38, CD138, GL-7, CD21, CD23, CD93 (AA4.1), CD11b, CD5, IgM and CD44 (eBioscience) for 30 minutes at 4°C in FACS staining buffer (PBS 1x with 5% FCS). Analysis and cell sort were then performed on a MoFlo instrument (Dako-Cytomation). B cell populations were defined as follows: GC (B220+ CD138-CD38lo/-GL-7+) and FO (B220+ CD138-CD21lo/neg CD23+ GL-7-). Cells were collected directly in sterile tubes containing supplemented OptiMEM (GIBCO) for cell culture.

### GC B cell culture in limiting dilution assay (LDA)

Decreasing number of GC B cells were cultured in 250 μl of OptiMEM (GIBCO) supplemented with 10% heat-inactivated FBS (GIBCO), 2 mM L-glutamine, 1 mM sodium pyruvate, 50 μM 2-ME, 100 U penicillin, and 100 μg/ml streptomycin. All cultures were performed in 96-well flat bottom plates containing 3×10^3^ NB40L feeder cells/well as previously described protocol with minor modifications [80]. GC B cells were added starting from 3000 cells/well to 1 cell/well through 3-fold dilution steps in the presence of 30 μg/ml of LPS (Salmonella typhimurium, Sigma-Aldrich) and 2 ng/ml of IL-21 (Peprotech). After 7 days, cultures were screened by ELISA to determine the frequency of IgG secreting GC B cells according to the Poisson distribution [81, 82].

### Single GC B cell cultures

Single GC B cells were sorted into 96-well round-bottom plates containing 3×10^3^ NB21 cells/well as previously described with minor modifications, mostly not adding IL-4 [54]. Supernatants were collected after 7 days of culture and cells were frozen in TCL lysis buffer supplemented with 1%β-mercaptoethanol for *Igh* sequencing.

### Igh sequencing

Single GC B cells from popliteal lymph nodes of Balb/c mice immunized with ZIKV or DENV NS1 proteins were sorted into 96-well PCR plates directly or after 7 days in culture. Plates contained 5 µl of TCL lysis buffer (Qiagen) supplemented with 1%β-mercaptoethanol. RNA extraction was performed using SPRI bead as described in [73]. RNA was reverse transcribed into cDNA using an oligo (dT) primer. *Igh* transcripts were amplified as described in [83]. PCR products were barcoded and sequenced utilizing MiSeq (Illumina) Nano kit v.2 as described in [84].

Paired-end sequences were assembled with PandaSeq [85] and processed with the FASTX toolkit. The resulting demultiplexed and collapsed reads were assigned to wells according to barcodes. High-count sequences for every single cell/well were analyzed. Ig heavy chains were aligned to both IMGT [86] and Vbase2 [87] databases, in case of discrepancy IgBLAST was used. V_H_ mutation analyses were restricted to cells with productively rearranged *Igh* genes, as described in [84]. CDR-H3 analyses were performed as described in [88]. Average hydrophobicity of CDR-H3 was calculated as previously described in [89]. Functional rearrangements were grouped by clonotypes defined by the same V_H_ and J_H_ segment and identical CDR-H3 length and amino acid sequence.

### Statistical analyses

Statistical analyses were performed using GraphPadPrism 7.0 software. Tests were chosen according to the type of variable and indicated in each result. Results with p> 0.05 were considered significant.

Principal component analysis (PCA) was performed to compare the repertoires globally. Reactivity sections were defined and signal intensities across the sections were quantified and analyzed as described in [90].

## Acknowledgements

We thank Dr. Orlando Ferreira for kindly providing ZIKV and DENV EDIII recombinant proteins; Dr. Edgar F.O. de Jesus (*in memoriam*) and his lab members for cell irradiation and Dr. Garnett Kelsoe for kindly providing the NB21 feeder cells. We thank Dr. Marcelo Bozza for helpful discussion and suggestions. This work was supported by the Brazilian research funding agencies Fundação de Amparo à Pesquisa do Estado do Rio de Janeiro (FAPERJ), Conselho Nacional de Desenvolvimento Científico e Tecnológico (CNPq), Coordenação de Aperfeiçoamento de Pessoal de Nível Superior (CAPES). CBC was supported by CNPq (PhD fellowship), and CAPES (PDSE 88881.132337/2016-01 and Projeto de pesquisa 1759/2014 - Biocomputacional - process number 23038.004628/2014-66). G.D.V. is a Burroughs-Wellcome Investigator in the Pathogenesis of Infectious Disease.

## Supplemental material

**Figure S1.**
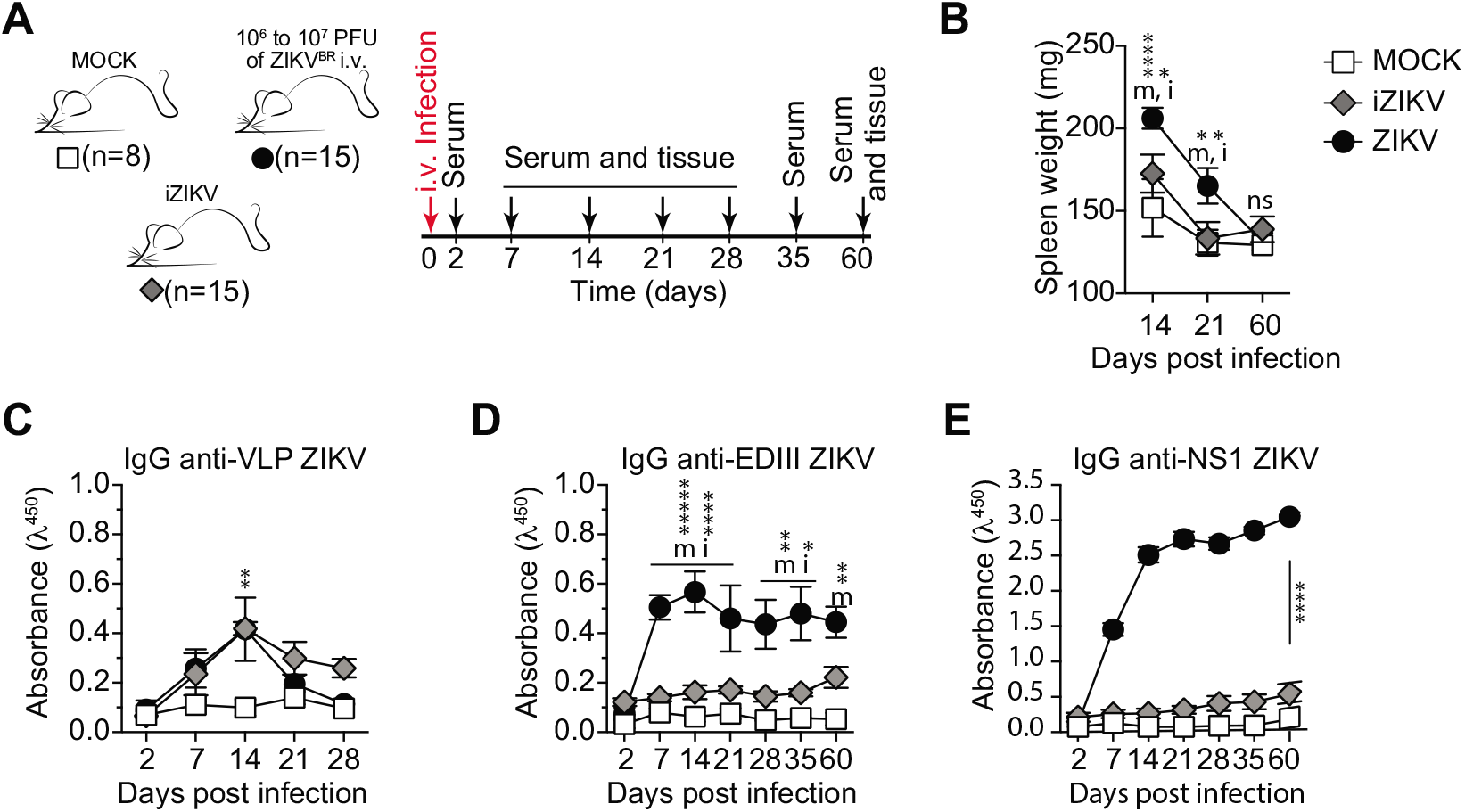
Characterization of humoral immune response to UV-inactivated ZIKV. **(A)** Experimental design indicating the time points of serum samples and lymphoid tissue collections from control mice (MOCK), mice immunized UV-inactivated virus (iZIKV) and infected mice (ZIKV). **(B)** Spleen weight measured at the time of collection, as indicated. (**C-E**) Serum levels of IgG specific to VLP (C), domain III of ZIKV envelope protein (D) and NS1 (E). Measured by ELISA at 1:120 dilution.

**Figure S2.**
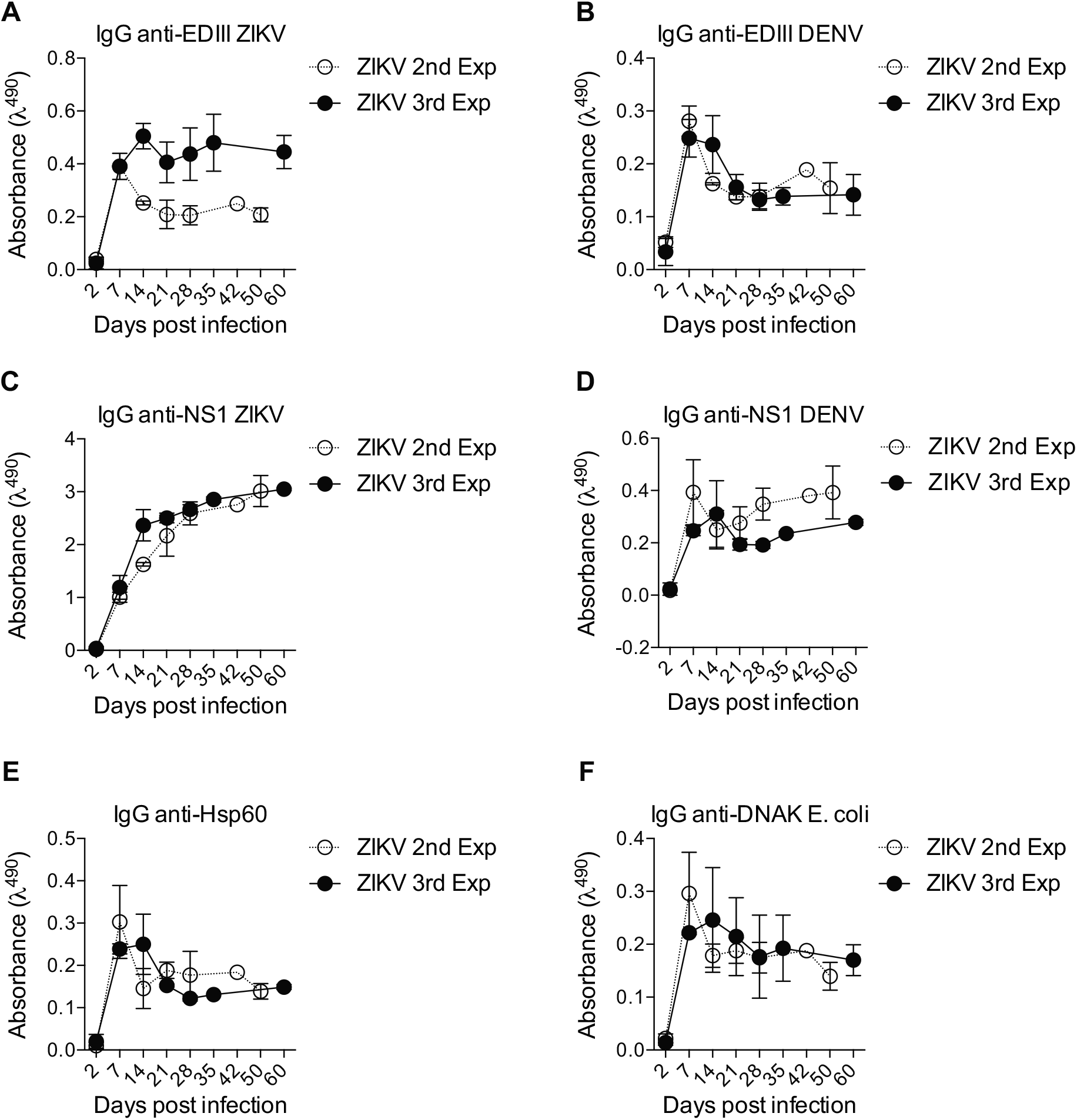
Cross reactivity with unrelated antigens. Sera from ZIKV infected mice (1:120 dilution) from 2 independent experiments were tested by ELISA for binding to ZIKV and DENV antigens EDIII **(A and B)** and NS1 **(C and D)** as well as unrelated antigens heat-shock protein Hsp60 and bacterial DNAK from E. coli at different time points. 2^nd^ experiment ended at 50 days and 3^rd^ experiment at 60 days.

**Figure S3.**
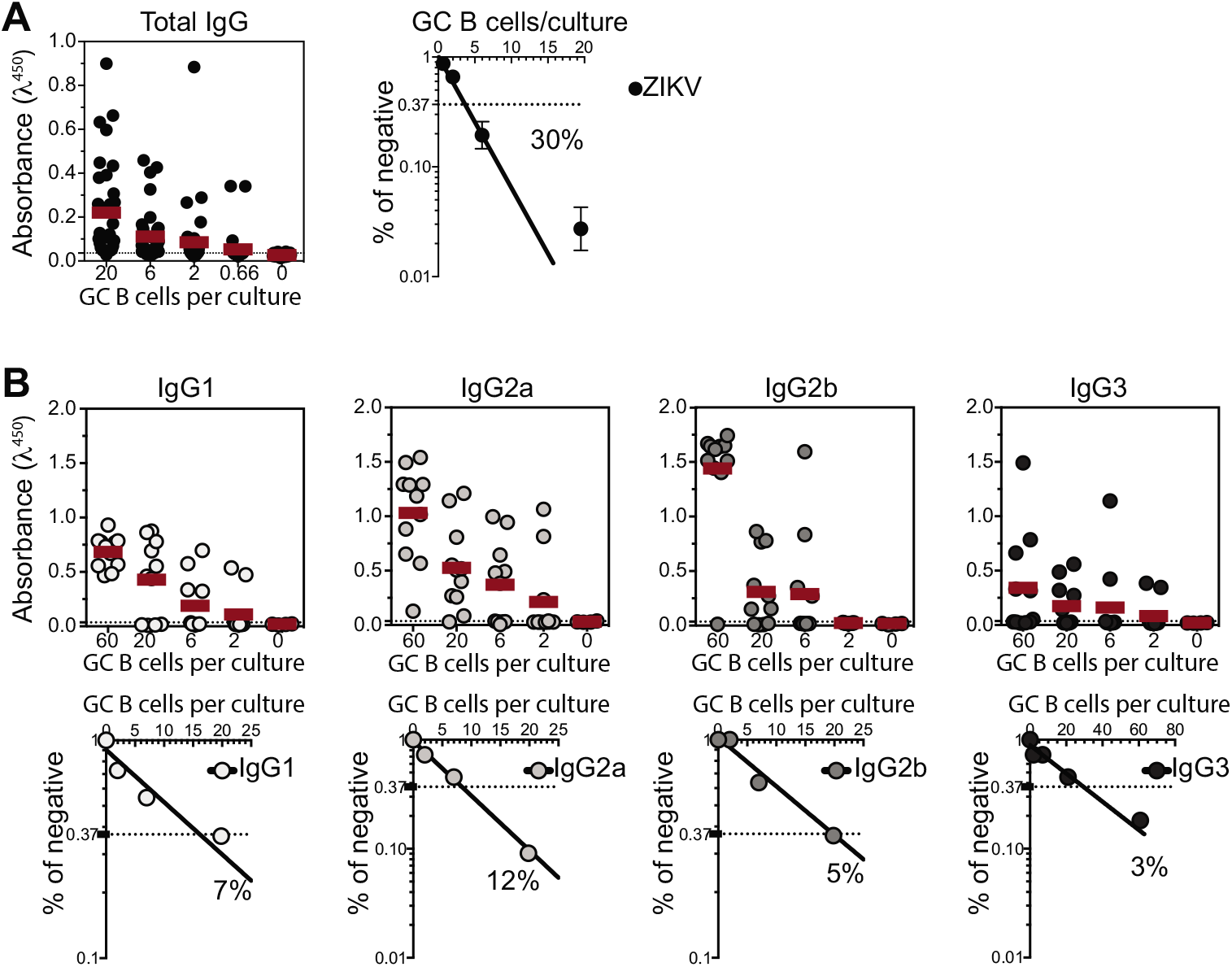
Quantification of the number of responding GC B cell clones per culture. GC B cells from popliteal lymph nodes of ZIKV-infected mice were sorted and cultured in decreasing average number of cells per well (60, 20, 6, 2, 0.66 cells/well). Supernatants were collected on day 7 and screened for IgG secretion by ELISA. **(A)** Frequency of GC B cell clones secreting total IgG per culture in response to polyclonal stimuli were calculated using *Poisson* distribution. **(B)** Culture supernatants were used to estimate the frequency of GC B cell clones secreting each BALB/c IgG subclasses (IgG1, IgG2a, IgG2b and IgG3). Cell culture was performed on a monolayer of gamma-irradiated (20 Gy) NB40L feeder cells (3 × 10^3^ cells/well), LPS (30 µg/mL) and IL-21 (2 ng/mL).

**Figure S4.**
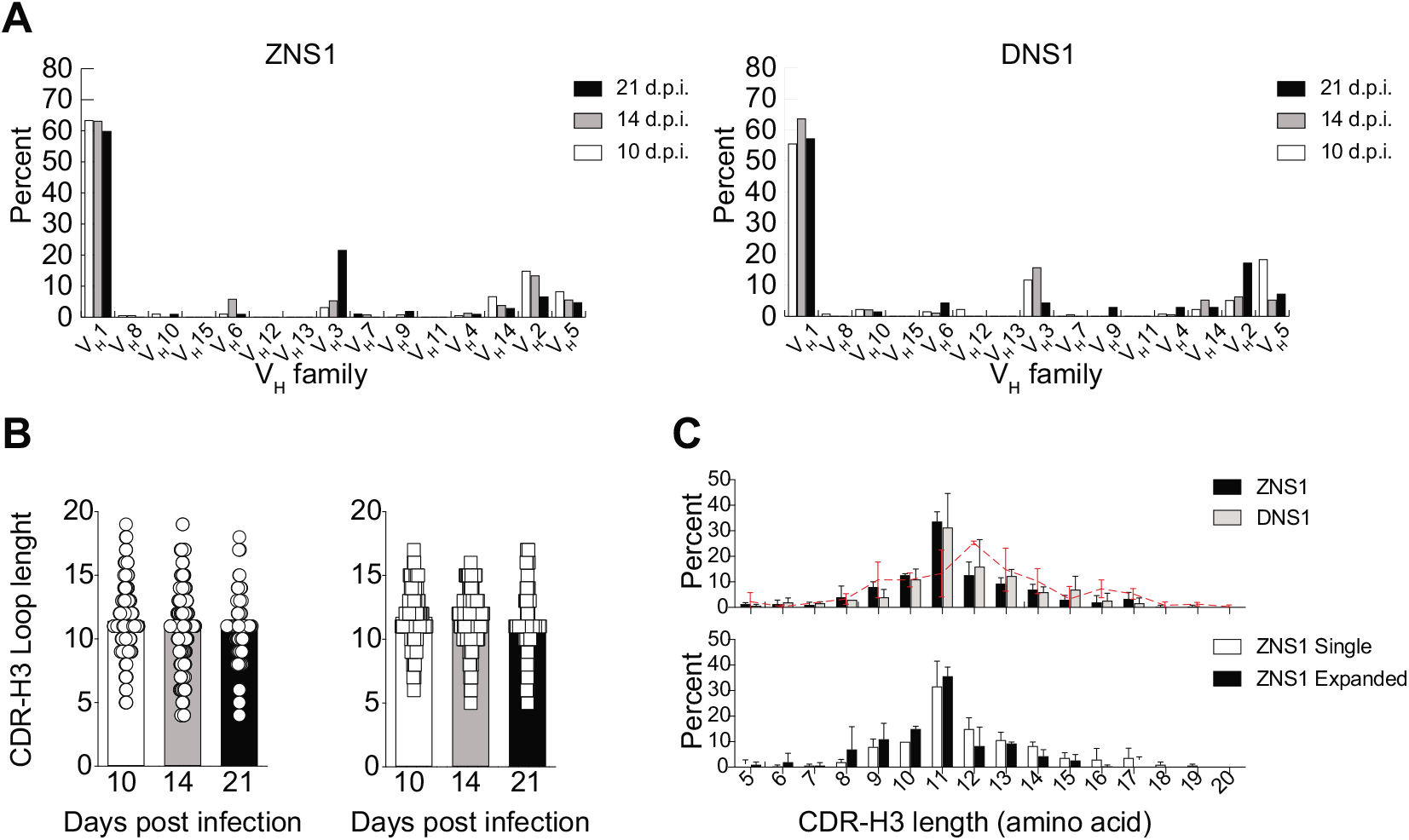
Analysis of V_H_ family usage and CDR-H3 length of *Igh* transcripts from B cells present in GCs after ZIKV NS1 or DENV NS1 immunization. **(A)** Shown here is the expression of each V_H_ family as a percentage of total functional transcripts from GC B cells at different time points after immunization with ZIKV NS1 or DENV NS1. **(B)** CDR-H3 loop length variation at different time points after immunization with ZIKV NS1 or DENV NS1. **(C)** CDR-H3 loop length distribution of all sequences at all time points (upper panel) and comparison between clonal (clonotypes found more than once in the same lymph node) and single clones from mice immunized with ZIKV NS1 (lower panel). Dashed red line indicates the distribution of CDR-H3 length in follicular B cells from WT BALB/c mice.

**Figure S5.**
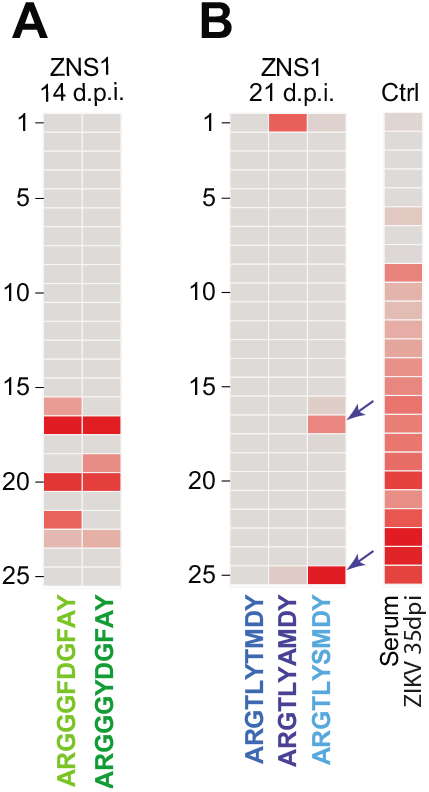
Heatmaps showing the pattern of self-reactivities of two particular clonal families. **(A)** Immunoreactivity profile of two poly-reactive variants from the clonal family bearing “glycine-enriched CDR-H3” (ARGGGYDGFAY) found among GC B cells from mice 14 d.p.i. with ZIKV NS1. **(B)** Immunoreactivity profile of the autoreactive clonotype ARGTLYAMDY and its two more mutated variants: ARGTLYTMDY (non-autoreactive) and ARGTLYSMDY (autoreactive).

**Supplementary Table 1:**
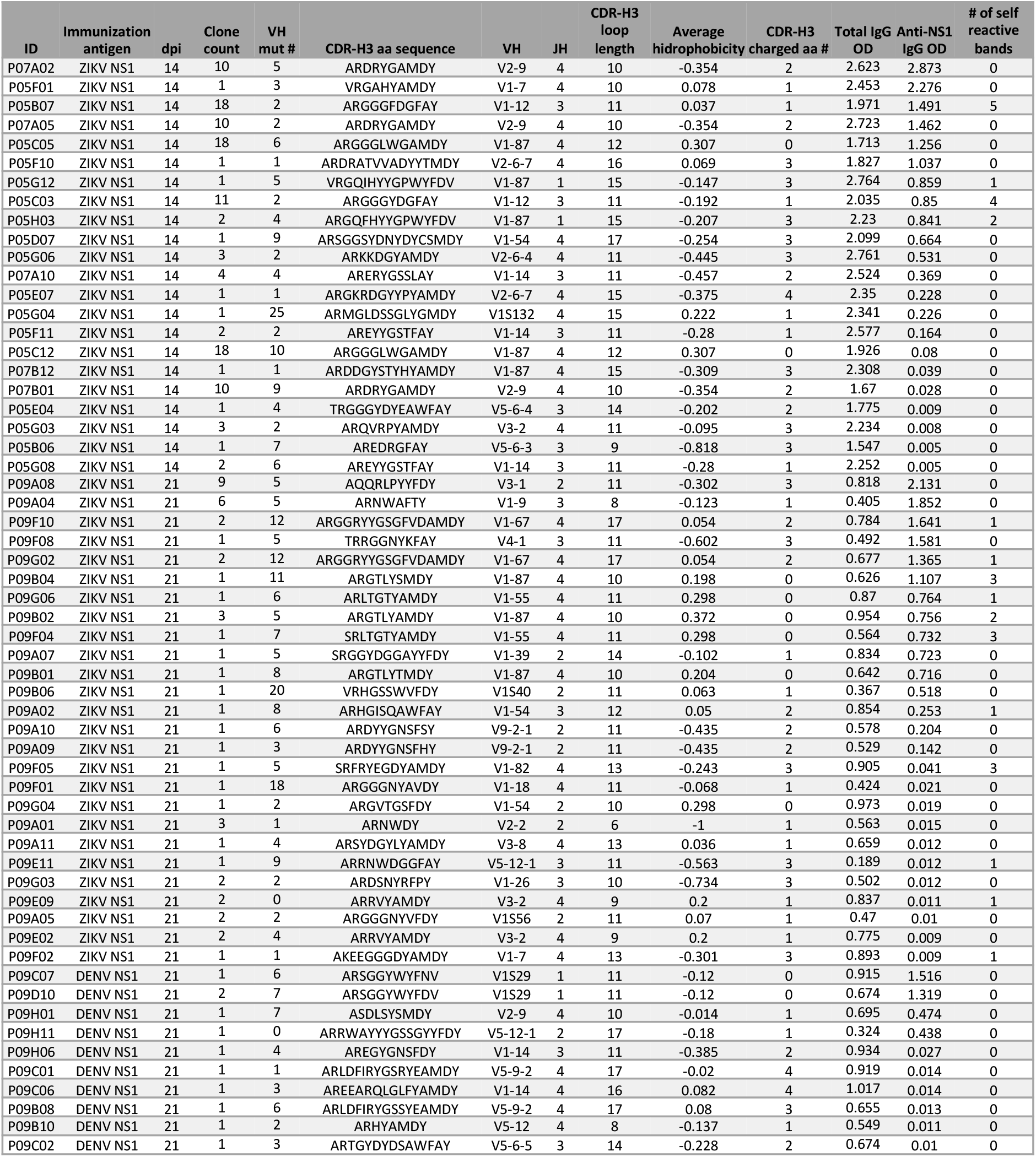
CDR-H3 characteristics from germinal center B cell clones tested for self-reactivity.

